# Correction for age-appropriate relaxation reference values alters estimates of neurometabolite concentration, but not age relationships

**DOI:** 10.64898/2026.11.23.768649

**Authors:** Yulu Song, Christopher W. Davies-Jenkins, Saipavitra Murali-Manohar, Kathleen E. Hupfeld, Gizeaddis L. Simegn, Steve C.N. Hui, Tao Gong, Vivek Yedavalli, Zahra Shams, Abdelrahman Gad, Aaron Gudmundson, Helge J. Zöllner, Georg Oeltzschner, Guangbin Wang, Eric C. Porges, Richard A.E. Edden

## Abstract

MRS-measured metabolite concentration estimates in the brain show significant changes over the lifespan. An important step in accurately quantifying metabolite levels using *in vivo* MRS is correction for T_1_- and T_2_-weighting of water and metabolite signals. Despite substantial literature indicating that both T_1_ and T_2_ of water and metabolites change across the lifespan, most aging studies use age-invariant reference values in quantification equations. It is important to establish whether observed changes in metabolite levels are driven by relaxation effects or actual concentration changes. In three recent studies, we have characterized changes in water and metabolite relaxation times across the adult lifespan, specifying linear models to predict age-appropriate T_1_ and T_2_ values for use in quantification. To establish whether prior metabolite-age correlation findings are driven by age-dependent errors in relaxation corrections, age-appropriate metabolite T_1_ and T_2_ values and age- and region-appropriate water T_1_ and T_2_ values were applied to re-calculate metabolite levels for a large lifespan cohort. While major metabolite (total N-acetyl aspartate tNAA, total choline tCho, total creatine tCr, *myo*-inositol mI, glutamate + glutamine Glx) concentration estimates were lower after using an age-appropriate relaxation correction analysis pipeline, the metabolite-age relationships were unchanged. We conclude that well-documented changes in metabolite and water relaxation times across the healthy adult lifespan do not drive findings of metabolite-age correlations.

## Introduction

Understanding neurometabolic changes associated with aging is critical for characterizing healthy brain aging^1–3^ and for investigating neurodegenerative disease^4,5^. Proton MRS is a non-invasive technology to measure neurometabolite levels, typically using the water signal as an internal reference^6–8^. Accurate concentration referencing requires compensation for the relaxation behavior of MR signals in brain tissue. MRS studies have long reported changes in neurometabolite levels across the lifespan^1–3^. T_1_-weighting usually arises in MRS experiments from the TR-dependence of the signal being excited (i.e., saturation-recovery). T_2_-weighting, on the other hand, arises from the TE-dependent exponential loss of signal between excitation and acquisition. Both relaxation processes arise from a complex interaction of molecular motion and interaction with the magnetic microenvironment, and both T_1_ and T_2_ are different for different metabolites. More recently, it has been shown that metabolite and water relaxation times also change with age^9–13^.

Physiologically, brain aging is associated with neuronal shrinkage, dendritic and synaptic loss, demyelination, metabolic slowing^14^, and microglial activation^15^. These physiological changes are associated with a number of distinct effects that can influence relaxation: changes in water content, metal deposition and demyelination. In healthy adults, gray matter (GM) water content tends to decrease with age, whereas white matter (WM) water content increases with age^16^. Changes in tissue water content and microstructure with aging alter T_1_ and T_2_ relaxation times. In parallel, iron accumulates in deep GM with aging^17^. Excess accumulation of iron can contribute to neurodegeneration and myelin breakdown, as observed in Alzheimer’s disease^18^. The deposition of iron and manganese also alters the T_1_ and T_2_ relaxation times of the tissue^19,20^. Myelin breakdown reduces the macromolecular content of tissue, which increases both T_1_ and T_2_ relaxation times^21^.

Relative changes in metabolites signal amplitudes across the lifespan may be driven by changes in metabolite concentration, but they may also be driven by differences in metabolite-specific relaxation times with age – a confounding factor that is largely unaddressed in the literature. Indeed, the majority of studies perform “relaxation correction”, but with reference parameters that are age-invariant, ignoring known changes in relaxation times across the lifespan^22–27^. Our ability to demonstrate significant changes in metabolite relaxation experimentally is limited by signal resolution, signal-to-noise ratio (SNR), cohort size, and statistical power. The age-dependence of T_1_ relaxation times has been shown for NAA in the centrum semiovale (CSO)^6,11,28^ and NAA and tCr in the posterior cingulate cortex (PCC)^11^. The age-dependence of T_2_^10,24,29–31^ has been shown for more metabolites, with additional methodological complexity associated with TE-modulation due to *J*-coupling. It is likely that relaxation behavior changes across the lifespan for all metabolites, to some degree, since the significant results reported tend to cluster around the most prominent signals. The relaxation behavior of tissue water, the most widely used MRS reference signal^6–8^, is also age-dependent^9^. Community consensus highlights the importance of relaxation correction for accurately quantifying metabolite levels using *in vivo* MRS^8,32^; it is every bit as important to correct for the age-dependence of T_1_- and T_2_-weighting in studies of neurometabolite changes with age.

In three recent studies^9–11^, we have characterized changes in water and major metabolite relaxation times across the adult lifespan, with robust sample sizes, reporting linear models to allow for age-appropriate T_1_ and T_2_ quantification corrections. The purpose of this work is to apply these new reference models to a previously reported lifespan cohort^3^, to *re-calculate* metabolite levels using age-appropriate relaxation reference values, and to test the extent to which prior findings were driven by age-dependence in relaxation (or age-dependence in the errors associated with relaxation correction).

## Methods

### Participants

102 healthy volunteers were recruited with local IRB approval. The adult cohort was structured to include approximately equal numbers of male and female participants in each decade of age from the 20s, 30s, 40s, 50s, to the 60s^3^. Exclusion criteria included contraindications for MRI and a history of neurological and psychiatric illness. Note that this cohort of subjects has been previously reported^3^ to investigate the age trajectory of metabolite levels (using age-invariant relaxation corrections). These data are publicly available: https://www.nitrc.org/projects/mm_mrs. Additional details of the cohort and protocols can be found in reference^3^.

### Data Acquisition Protocol

Data were acquired on a 3T MRI scanner (Ingenia CX, Philips Healthcare, The Netherlands). Acquisition of MRS data was preceded by a T_1_-weighted MP-RAGE scan (TR/TE 6.9/3.2 ms; FA 8°) with 1 mm^3^ isotropic resolution for voxel positioning and tissue segmentation. Metabolite spectra were acquired using PRESS localization with the following parameters: TR/TE 2000/30 ms; 30 x 26 x 26 mm^3^ voxels localized in the CSO (predominantly WM) and PCC (predominantly GM), as shown in Figure 1; 96 transients sampled at 2 kHz; water suppression was performed using the VAPOR^33^ method. More details of this dataset can be found in previous published paper^3^.

**Figure 1.**
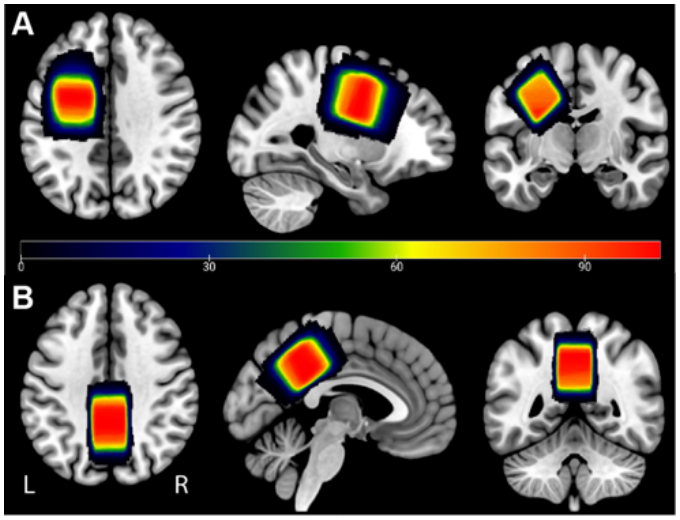
Cohort voxel placement. MRS data were acquired from **A** CSO and **B** PCC. Each participant’s native space binary voxel masks were normalized to standard space (Montreal Neurological Institute MNI) and overlaid. Warmer colors indicate areas of greater overlap between participants (color bar shows number of subjects overlapped).

### MRS Data Analysis

Osprey 2.9.0^34^, an open-source analysis toolbox for MRS data, was used to process PRESS data within MATLAB R2024a. The analysis procedures followed consensus-recommended processing guidelines for MRS data^8,32^. Briefly, standardized analysis steps in Osprey included eddy-current correction based on the unsuppressed water reference^35^, frequency- and phase-correction using probabilistic spectral alignment^36^, removal of residual water signals using a Hankel singular value decomposition (HSVD) filter^37^, and modeling of the metabolite peaks^34,38^. An appropriate custom basis set was generated using MRSCloud^39^ with simulated profiles for following metabolites: ascorbate (Asc), aspartate (Asp), creatine (Cr), methylene protons of Cr (CrCH_2_), *γ*-aminobutyric acide (GABA), glycerophosphocholine (GPC), glutathione (GSH), glutamine (Gln), glutamate (Glu), *myo*-inositol (mI), Lactate (Lac), NAA, N-acetyl aspartyl glutamate (NAAG), phosphocholine (PCh), phosphocreatine (PCr), phosphoryl ethanolamine (PE), scyllo-inositol (sI), taurine (Tau), parameterized macromolecules (MMs)^40^, Lip09, Lip13, and Lip20. The model spectral range was from 0.5 to 4.0 ppm with a default Osprey baseline knot spacing of 0.4 ppm. Amplitude-ratio soft constraints were applied to the amplitudes of the lipids, and NAAG/NAA peaks as defined in the LCModel manual (http://s-provencher.com/pub/LCModel/manual/manual.pdf). The water-reference data were modeled in the frequency domain with a dedicated water basis function and a six-parameter model (amplitude, zero- and first-order phase, Gaussian and Lorentzian line broadening, and frequency shift). More details are in Supplementary Table 1 MRSinMRS.

### Quantification

Metabolite concentration estimates were derived from linear-combination modeling coefficients according to the well-established Gasparovic^41^ tissue- and relaxation-correction model (Equations [1] and [2]): once with the legacy age-invariant constant reference values for water and metabolite T_1_ and T_2_ (i.e., the previous default relaxation values used by Osprey, taken from the literature^7,12,13^), as shown in blue in Table 1; and once with their age-appropriate reference values calculated from the published linear models^9–11^, as shown in red in Table 1. For comparison, results from the original analysis^3^, were rerun with an updated version of Osprey (Osprey 2.9.0) and are also presented, as shown in black in Table 1. The key distinction between the original analysis pipeline (in black) and the water-atlas analysis pipeline (in blue) lies in the application of region-appropriate water reference values that are calculated from the water relaxation atlas we synthesized previously^9^,

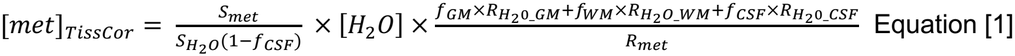

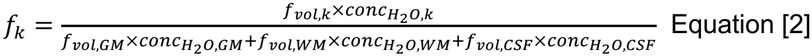

where *S*_*met*_ is the metabolite signal amplitudes; [H_2_O] is the molal concentration of pure MR-visible water, i.e., 55.5 mol/kg of MR-visible water^41,42^; *f*_*GM*_, *f*_*WM*_, *f*_*CSF*_ are the molal water fractions for GM, WM and CSF, which are derived from the volume fractions according to Equation [2], with the relative water densities 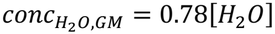,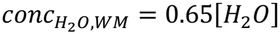 and 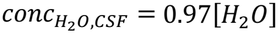. Tissue-specific relaxation corrections are calculated according to [3] 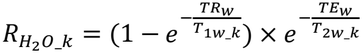 and [4] 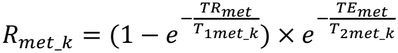, where *TR*_*w*_, *TE*_*w*_, *TR*_*met*_, *TE*_*met*_ are the repetition and echo times of the water-unsuppressed and water-suppressed acquisitions; *T*_1*w*_*k*_ and *T*_2*w*_*k*_ are tissue-specific relaxation times for water; and *T*_1*met*_*k*_ and *T*_2*met*_*k*_ are the tissue-specific relaxation times of metabolites.

**Table 1.** Specification of the three analysis pipelines.

|  | Water region-dependent | Water age-dependent | Water values reference | Metabolite Age-dependent | Metabolite references |
|---|---|---|---|---|---|
| original analysis | no | no | 7 | no | 12,13 |
| water-atlas | yes | no | 9 | no | 12,13 |
| age-appropriate | yes | yes | 9 | yes | 10,11 |

Metabolite relaxation models are summarized in Figure 2. Metabolite concentration estimates, calculated with the three analysis pipelines, were plotted against age in Figure 3. We also explored the predicted relaxation weighting that affects metabolite signals at different TEs and TRs against age in Figure 4.

**Figure 2.**
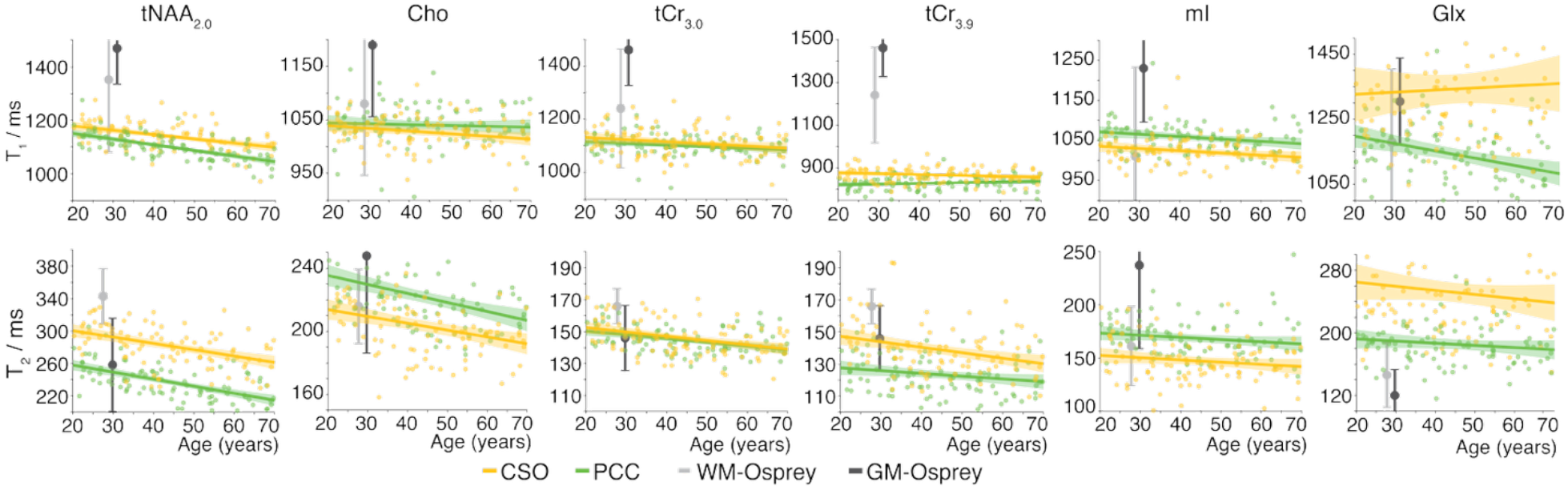
Linear models of metabolite T_1_ and T_2_ values as a function of age are represented for the CSO voxel (gold) and the PCC voxel (green). The shaded area represents the 95% confidence intervals for the linear models^10,11^. These plots synthesize previously published data applied here and Osprey default values, shown as single points for GM (dark gray) and WM (light gray), with whiskers representing the reported standard deviation^12,13^. Note, the T_1_ default values were derived from a cohort with a mean age of 30 years^13^, and T_2_ values from a cohort with a mean age of 28.7 years^12^.

**Figure 3.**
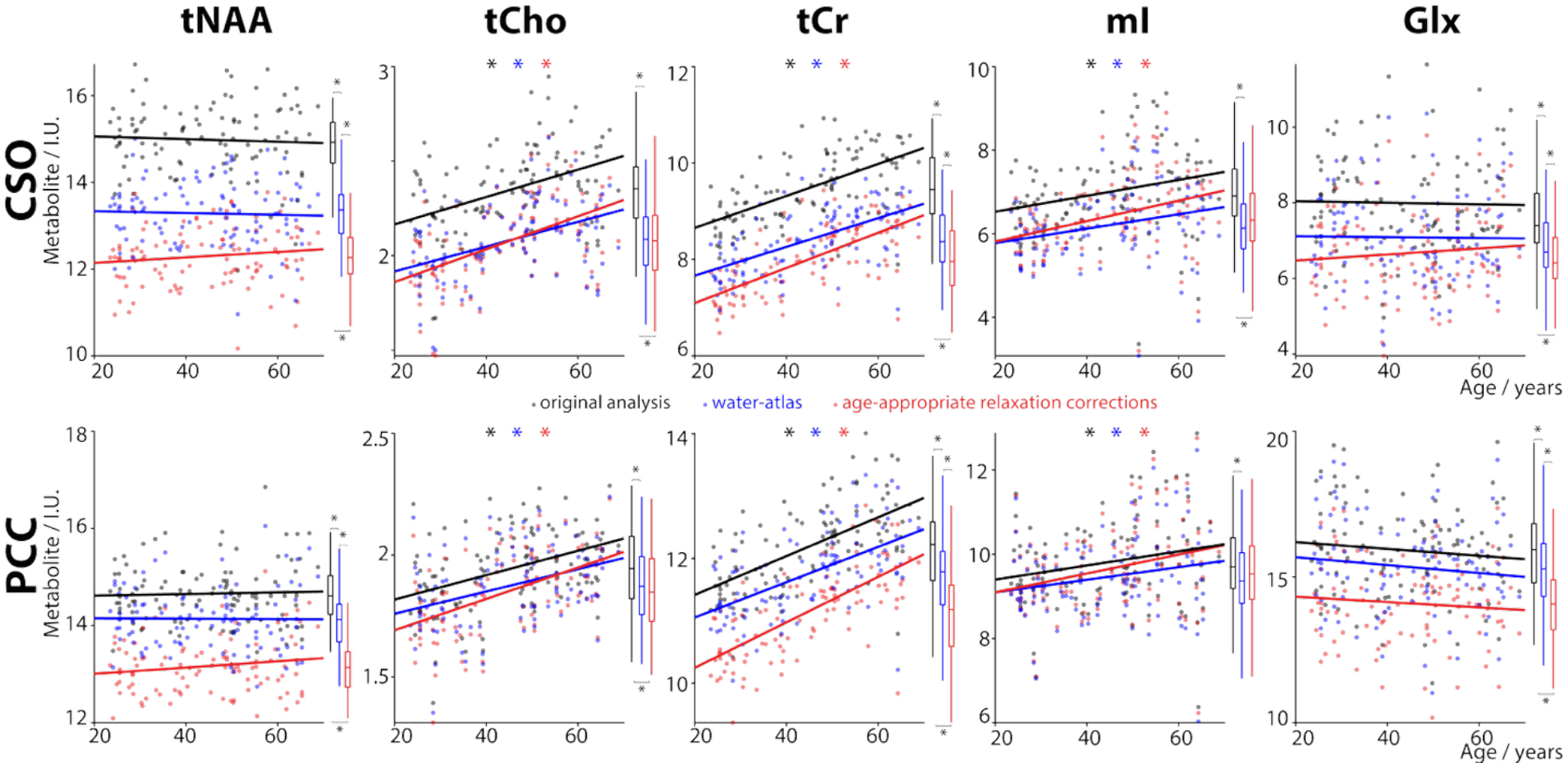
Metabolite-age correlations for CSO (top) and PCC (bottom) voxels, with overall distributions shown in the boxplots. Black datapoints repeated the original published analysis (but with a newer version of Osprey). Blue datapoints underwent relaxation correction with water-atlas reference values for metabolite and region-appropriate water relaxation values. Red datapoints underwent relaxation correction with age-appropriate reference values for metabolite and age- and region-appropriate water relaxation values, based upon recent findings. * *p* < 0.05.

**Figure 4.**
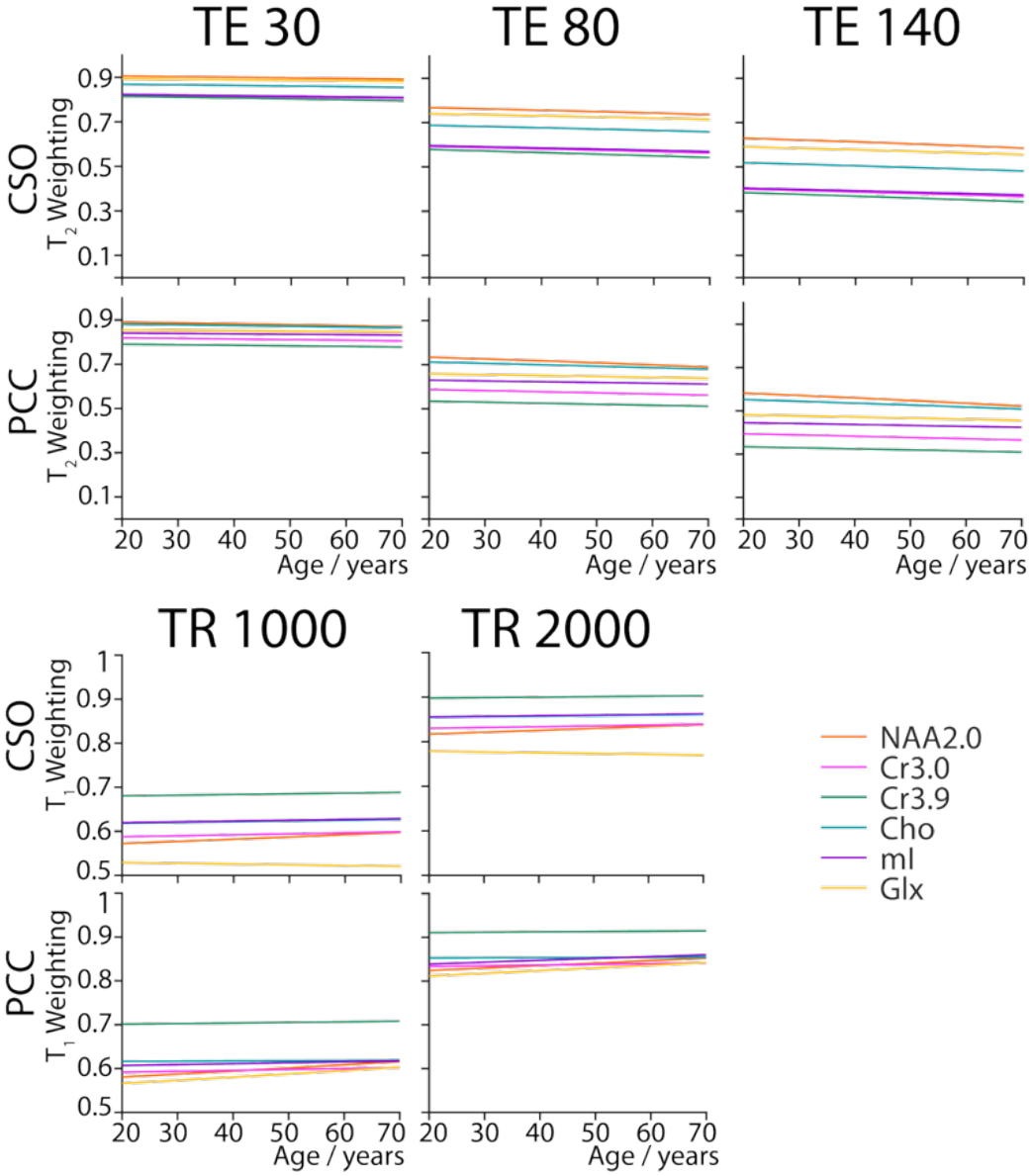
Predicted metabolite T_1_- and T_2_-weighting as a function of age. For typical acquisition ranges of TE and TR, relaxation weightings are less strongly age-dependent than the relaxation times themselves

### Statistical Analysis

All statistical analyses were performed using R (Version 4.5.0) in RStudio (Version 2024.12.1.563). Data were analyzed for: total NAA (tNAA = NAA + NAAG); total choline (tCho = GPC + PCh); total creatine (tCr = Cr +PCr); Glx = Glu + Gln; and mI. Concentration values of 0 were considered indicative of a fitting failure, and the corresponding data points were excluded from further analyses. An ANOVA test was utilized to compare the original analysis pipeline, the age-invariant water-atlas analysis pipeline and the age-appropriate relaxation correction analysis pipeline, to assess whether the means of the three analysis pipelines are significantly different. Prior to conducting the ANOVA test, the data were examined for normality (assessed using the Shapiro-Wilk test) and homogeneity of variances (assessed using Levene’s Test) to ensure the appropriateness of this parametric approach. Associations between age and metabolite concentrations were examined on a voxel-wise basis. For each voxel, a Pearson correlation was performed when both variables met the normality assumption (Shapiro-Wilk test, p > 0.05). If either distribution deviated from normality, a Spearman rank-order correlation was used instead. Two-one-sided equivalence tests (TOST)^43^ were carried out to determine whether the observed correlation were statistically equivalent to zero, using a predefined equivalence interval of |r| < 0.30.

## Results

The demographic characteristics of 102 participants are presented in Table 2. This is a well-balanced (age and gender) dataset, making it an appropriate cohort to use to investigate the age-relaxation confound in MRS studies of aging.

**Table 2.** Demographic characteristics of participants.

|  | 20s | 30s | 40s | 50s | 60s | Total |
|---|---|---|---|---|---|---|
| Age (years) |  |  |  |  |  |  |
| mean | 26.0 | 34.6 | 45.7 | 53.9 | 63.6 | 44.5 |
| Sex, n |  |  |  |  |  |  |
| Male | 10 | 10 | 10 | 10 | 9 | 49 |
| Female | 11 | 11 | 10 | 10 | 11 | 53 |
| Total, n | 21 | 21 | 20 | 20 | 20 | 102 |

**Table 3.**
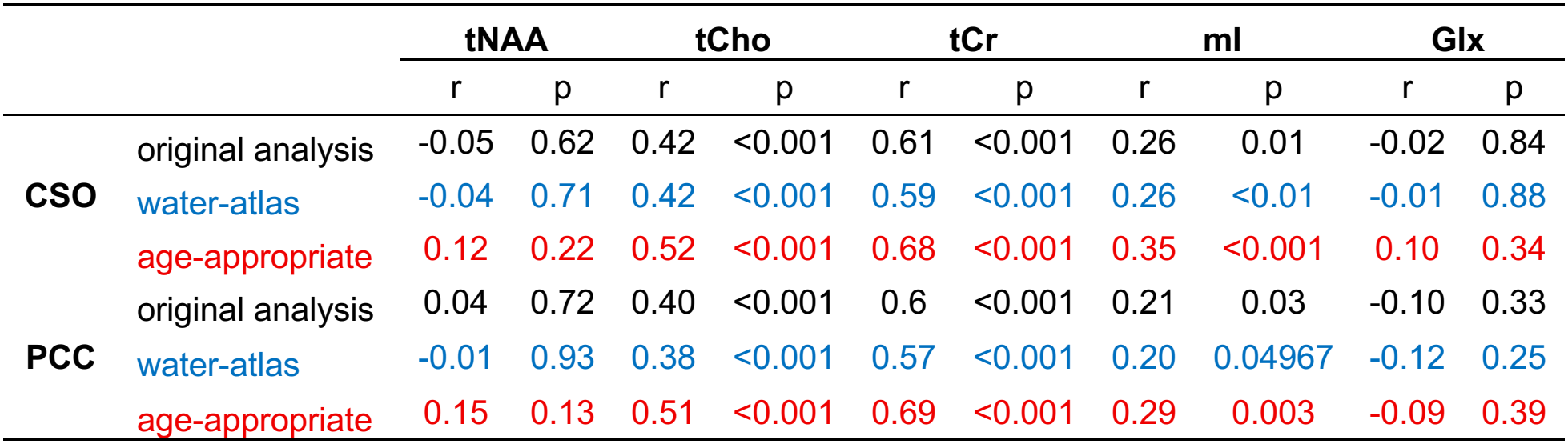
Correlation coefficients (r) and corresponding p values for each metabolite, region and analysis pipelines.

|  |  | **tNAA** |  | **tCho** |  | **tCr** |  | **mI** |  | **Glx** |  |
|  |  | r | p | r | p | r | p | r | p | r | p |
|---|---|---|---|---|---|---|---|---|---|---|---|
| **CSO** | original analysis | -0.05 | 0.62 | 0.42 | <0.001 | 0.61 | <0.001 | 0.26 | 0.01 | -0.02 | 0.84 |
|  | water-atlas | -0.04 | 0.71 | 0.42 | <0.001 | 0.59 | <0.001 | 0.26 | <0.01 | -0.01 | 0.88 |
|  | age-appropriate | 0.12 | 0.22 | 0.52 | <0.001 | 0.68 | <0.001 | 0.35 | <0.001 | 0.10 | 0.34 |
| **PCC** | original analysis | 0.04 | 0.72 | 0.40 | <0.001 | 0.6 | <0.001 | 0.21 | 0.03 | -0.10 | 0.33 |
|  | water-atlas | -0.01 | 0.93 | 0.38 | <0.001 | 0.57 | <0.001 | 0.20 | 0.04967 | -0.12 | 0.25 |
|  | age-appropriate | 0.15 | 0.13 | 0.51 | <0.001 | 0.69 | <0.001 | 0.29 | 0.003 | -0.09 | 0.39 |

### Metabolite levels

In the predominantly WM CSO region, one-way ANOVA results revealed a significant effect of relaxation correction method on tNAA, tCr, and Glx concentration estimates (p-value < 0.001 for both, effect size = 0.71 (tNAA), 0.44 (tCr), 0.17(Glx)). Post-hoc comparisons indicated that metabolite levels were significantly lower using the age-appropriate relaxation-corrections pipeline compared with both the original analysis pipeline (p < 0.001 for tNAA, tCr, and Glx), and the water-atlas analysis pipeline (p < 0.05 for tNAA, tCr, and Glx). Metabolite level estimates were also significantly lower using the water-atlas analysis pipeline compared with the original analysis pipeline (p < 0.001 for tNAA, tCr, and Glx).

One-way ANOVA results revealed a significant effect of relaxation correction method on tCho and mI levels (p-value < 0.001 for both, effect size = 0.25 (tCho), 0.13(mI)). Post-hoc comparisons indicated that metabolite levels were significantly lower using the age-appropriate relaxation-correction pipeline compared with the original analysis pipeline (p < 0.001 for both tCho and mI). Metabolite level estimates were also significantly lower using the water-atlas analysis pipeline compared with the original analysis pipeline (p < 0.001 for both tCho and mI)).

In the predominantly GM PCC region, one-way ANOVA results revealed a significant effect of relaxation correction method on tNAA, tCr, and Glx levels (p-value < 0.001 for all, effect size = 0.51 (tNAA), 0.28 (tCr), 0.20 (Glx)). Post-hoc comparisons indicated that metabolite level estimates were significantly lower using the age-appropriate relaxation-correction pipeline compared with both the original analysis pipeline (p < 0.001 for tNAA, tCr, and Glx), and the water-atlas analysis pipeline (p

< 0.001 for tNAA, tCr, and Glx). Metabolite estimates were also significantly lower using the water-atlas analysis pipeline compared with the original analysis pipeline (p < 0.05 for tNAA, tCr, and Glx). One-way ANOVA results revealed a significant effect of relaxation correction method on tCho levels (p-value < 0.001, effect size = 0.05). Post-hoc comparisons indicated that tCho level estimates were significantly lower using the age-appropriate relaxation-correction pipeline compared with the original analysis pipeline (p < 0.001), and tCho level was also significantly lower using the water-atlas analysis pipeline compared with the original analysis pipeline (p < 0.01). One-way ANOVA results revealed a non-significant effect of relaxation correction method on mI levels (p-value = 0.06).

### Correlation Results

Correlation plots for levels of the major MRS signals (tNAA, tCho, tCr, mI, Glx), calculated using original analysis (in black), water-atlas (in blue) and age-appropriate relaxation corrections (in red), are shown in Figure 3. In both CSO and PCC regions, across the three analysis pipelines, tCho, tCr and mI levels exhibited a positive correlation with age (tCho p < 0.001 for all; tCr p < 0.001 for all; mI p < 0.05 for all), and age slopes tend to be slightly more positive in the age-appropriate relaxation-correction pipeline. Across all three analysis pipelines and in both brain regions, no significant correlations were found between age and the metabolite tNAA and Glx (all p >0.05). TOST (equivalence bounds ± 0.30) indicated that the observed correlations were statistically equivalent to zero (all p_TOST < 0.05), except tNAA using age-appropriate relaxation corrections (in red) in both CSO and PCC regions, where traditional test was non-significant and the TOST was also non-significant, indicating that the data are inconclusive: we have insufficient evidence to determine whether the true correlation is meaningfully different from zero.

### Prediction Analysis

The predictions of metabolite relaxation weighting at three different TEs (TE = 30 ms, 80 ms, 140 ms) and two TRs (TR = 1000 ms, 2000 ms) were plotted against age (as shown in Figure 4). With increasing TE, T_2_ weightings become more pronounced in both intercept and slope, as expected, for both CSO and PCC regions. The T_2_-weighting difference between different metabolites, as expected, also increases at longer TE. With increasing TR, T_1_ weighting decreased in intercept, but there is a less pronounced impact on slope. The T_1_-weighting difference between different metabolites decreases with increasing TR.

## Discussion

Relaxation correction is a crucial quantification step in improving the accuracy of MRS studies. Because a full subject-by-subject relaxometric work-up is generally impractical, most studies use literature-derived metabolite and water T_1_s and T_2_s to calculate the necessary corrections. This common practice, when it fails to accommodate age-related changes in relaxation behavior, introduces the potential to confound age-related changes in metabolite relaxation with changes in metabolite concentration. In this study, we implemented age- and region-appropriate relaxation correction for the major metabolites and water. We then investigated the practical impact of this procedure by conducting a re-analysis of a large healthy aging cohort. We found that metabolite concentration estimates were mostly significantly lower when age-appropriate relaxation corrections were applied. However, the correlations between MRS-measured metabolite levels and age did *not* change significantly. This suggests that previously reported metabolite-age correlations are likely *not* driven by relaxation time changes, as previously suspected, and do indeed reflect actual changes in metabolite concentrations.

Several factors contribute to the observed differences in metabolite levels. Firstly, changes in metabolite T_1_ and T_2_ relaxation times between ages 20 and 70 are relatively modest (Figure 2). Therefore, the application of age-appropriate relaxation times is expected to result in only minor adjustments in estimated metabolite levels across this age range. Conversely, the age-appropriate relaxation times differ substantially from the default reference values published in the literature, particularly for metabolite T_1_ values (Figure 2). Estimated metabolite levels are significantly lower with the age-appropriate analysis pipeline than with the original and age-invariant analysis pipelines, for all metabolites in CSO and PCC regions, except mI in PCC. Secondly, MRS acquisition parameters are typically selected to maximize SNR for accurate metabolite quantification. This involves minimizing T_1_- and T_2_-weighting—here, a relatively short TE of 30 ms, and a TR of 2 s (Figure 4). This reduces the effects of differential relaxation rates on metabolite signal intensities.

Other groups have argued that using even longer TR (~5 seconds) can virtually eliminate all T_1_ weighting without incurring much of an SNR penalty^44^, and may therefore be preferable over medium TRs (~2 seconds). Thirdly, age-related relaxation changes to the water and metabolite signals tend to be in the same direction, and when applied to the numerator and denominator respectively, tend to mitigate each other. Finally, the observed age-related decreases in T_1_ and T_2_ have opposing effects on estimated metabolite concentrations after relaxation correction (Equation [4]), thereby partially offsetting each other.

So, if the use of age-appropriate relaxation times has little impact on correlation findings, what is the value in making these measurements? Our well-powered findings directly establish this premise: at 3T, using short TE and relatively long TR, the correlations between MRS-measured metabolites and age remain largely unchanged after applying age-appropriate relaxation time corrections. This finding is, itself, useful for the interpretation of MRS studies of healthy aging, as it provides greater confidence that observed age-related changes in metabolite estimates are not substantially driven by the use of age-invariant relation times. In studies employing shorter TR (e.g., MRSI) or longer TE (e.g., edited MRS), relaxation weighting will be larger, and age-appropriate relaxation corrections will therefore have a more pronounced effect. The approach (and reference curves) applied here can be applied to directly assess the degree of the relaxation confound for other studies retrospectively.

Furthermore, the use of more appropriate (and more rigorously measured) reference values for relaxation is likely to results in more accurate reference values for age-normed metabolite concentrations. Although the slope and significance of metabolite-age correlations is largely unchanged between the three quantification pipelines compare, there is frequently a ‘vertical offset’ between results. These arise from mean-shifts in the reference values, as opposed to age effects which tend to mitigate each other.

For metabolite T_1_ relaxation values, the age-dependent relaxation models were obtained from Murali-Manohar et al^11^. The Osprey default values were derived from Mlynárik et al^13^, which was, at the time of designing Osprey, the most widely cited reference for metabolite T_1_ at 3T. On average the Mlynárik T_1_s are longer, resulting in stronger relaxation corrections, and are reported with relatively wide error estimates. Several methodological differences may contribute to the discrepancy between these reference values. First, the study populations differed substantially.

Murali-Manohar et al. aimed to characterize age dependency in neurometabolite T_1_ relaxation times by recruiting 101 participants across the adult lifespan. In contrast, Mlynárik et al. focused primarily on establishing a single reference T_1_ value, with less emphasis on age as a variable; their cohort consisted of only five participants aged 20-48 years. Second, the sequence and protocol differed.

Murali-Manohar et al. employed PRESS (TE/TR 30/2000 ms) with a relatively large voxel (30×26×26 mm^3^), whereas Mlynárik et al. used STEAM (TR 6 s) with a smaller voxel (20×20×20 mm^3^). Although the longer TR used by Mlynárik et al. allows nearly full recovery of longitudinal magnetization and therefore minimizes T_1_-weighting, it also increases scan duration, which may reduce temporal SNR or potentially increase motion artifacts. Smaller voxels reduce partial volume effects, allowing more precise measurement at the cost of less signal. Murali-Manohar et al. sampled eight TIs (300; 400; 511; 637; 780; 947; 1148 and 1400 ms) with finer steps, enabling denser and more precise modeling of the relaxation curve, whereas Mlynárik et al. used five to six TIs (150 – 1650 ms) with larger intervals (300 ms). Mlynarik quantified spectra by manually reading “peak heights”, wheras Murali-Manohar applied more sophisticated linear combination modeling. These differences in population, acquisition protocol, voxel size, relaxation sampling, and analysis could each contribute to differences in the resulting T_1_ estimates.

Similar considerations apply to metabolite T_2_ differences. The age-dependent T_2_ models were obtained from Hupfeld et al.^10^. The Osprey default values were derived from Wyss et al.^12^, which were on average longer (with a less clear distinction than for T_1_), and also reported with relatively wide error estimates. The study populations differed substantially: Wyss et al. included a predominance of young adults with an unequal sex distribution; whereas Hupfeld et al. analyzed the same cohort as Murali-Manohar et al., comprising a balanced sample across age and sex. The sequence and protocol differed: Hupfeld et al. employed conventional PRESS with comparably few echo times (8 steps ranging from 30 – 350 ms) and a TR of 2 s; in contrast, Wyss et al. used *J*-PRESS with one hundred 2-ms steps spanning a narrower range (30 – 228 ms), a shorter TR of 1.6 s and a post-hoc correction scheme for retrospective correction of T_2_. Hupfeld et al. employed a larger voxel (30×26×26 mm^3^) than Wyss et al. (25×20×20 mm^3^). Thus, differences in both the sampled populations and acquisition methodology may contribute to the different T_2_ estimates used by those analysis pipelines.

Three quantification methodologies were applied to disentangle different contributions to changes in metabolite measurements. The water-atlas methodology (blue) differs from the original analysis (black) in using region-appropriate reference values from Simegn et al.^9^, whereas the Osprey default values were derived from Wansapura et al.^7^. Overall, the water-atlas analysis accounted for the majority of the difference between the original analysis and the full age-dependent analysis (red).

This suggests that changes in the water reference values are most impactful, although in many cases, additional changes occurred between the intermediate and full age-dependent analysis, in the same direction. Wansapura et al. recruited 19 subjects aged 29 - 47 years, whereas the Simegn values were measured in the same age- and sex-balanced healthy control cohort as the metabolite models. Acquisition methods differed substantially: Simegn et al. used the DESPOT^45^ method, a comparatively time-efficient protocol, whereas Wansapura et al. employed a saturation-recovery acquisition to estimate T_1_ and a multi-spin-echo sequence for T_2_. It is well recognized that T_1_ and T_2_ measurements vary substantially with sequence design, acquisition sampling scheme, reconstruction algorithms, magnetization-transfer effects, and subject motion^46–49^. In the study by Simegn et al., relaxation-weighted water signals were first estimated for each microvoxel and averaged to create a whole-brain relaxation atlas. The water-atlas relaxation values differed from the defaults across tissue types: in GM, T_1_ increased from 1.3s to 1.9s in CSO and 1.8s in PCC, and T_2_ increased from 0.11s to 0.24s in CSO and 0.21s in PCC; in WM, T_1_ increased from 0.8 to 1.2s in CSO and 1.3s in PCC, while T_2_ altered from 0.08s to 0.07s in CSO and 0.09s in PCC. These parameters result in both T_1_ and T_2_ weighting terms in the range 0.66 to 0.91 for TR 2s and TE 30ms. The changes in T_1_ reference values result in a ~10% reduction across the board (i.e. for WM and GM, and for CSO and PCC). For CSO, the T_2_ of the major component (WM) does not change greatly, so this is the only effect in play. For PCC, the T_2_ of the major component (GM) changes substantially, resulting in a ~10% change in the weighting term, that mitigates the T_1_-based change. Overall, therefore, the water atlas reference values result in a meaningful reduction in values in CSO, and a smaller reduction in PCC.

Differences between the full pipeline (red) and the water-atlas (blue) quantification methods involve metabolite relaxation differences and the additional consideration of age-dependent water values. Because the metabolite reference models have an intercept at age-30, differences at the age-30 point are entirely explained by changes in the metabolite relaxation time intercepts. Across both regions, NAA, Cr and Glx saw a downward shift of ~8%, while tCho and mI were relatively unchanged. NAA and Cr shifts were mainly caused by the new T_1_ values, while Glx was driven by T_2_ corrections. tCho and mI tended to have smaller changes in relaxation weighting, and opposing shifts in T_1_ and T_2_ effects. The full pipeline model consistently showed a slightly more positive slope, for all metabolites and regions, driven principally by increases in the water GM T_2_ with age.

There are some limitations to this study. First, the age-specific metabolite relaxation values were derived only at 3T, covered two brain regions and five major metabolites across the adult lifespan, whereas the water-relaxation atlas covered the whole brain. Future work should expand the metabolite values into atlases, include additional metabolites, magnetic field strengths, and age range. Second, the water-relaxation atlas was derived using the DESPOT method rather than conventional multi-echo or inversion-recovery approaches for T_1_/T_2_ mapping. Differences in acquisition and modeling methodology may introduce systematic differences in the estimated relaxation times and therefore represent a potential source of uncertainty in the present analysis. Third, the cohort consisted exclusively of healthy controls. Although this provides an important baseline for understanding relaxation and metabolite changes in healthy aging, it may limit the generalizability of these findings to clinical populations.

## Conclusion

In conclusion, well-documented changes in metabolite and water relaxation times across the healthy adult lifespan do not drive findings of metabolite-age correlations, though accounting for these changes reduces estimated metabolite levels. After correction with age-appropriate relaxation reference values, prior metabolite-age correlation findings are largely unchanged.

## Acknowledgments

This work was supported by National Institutes of Health (NIH) grants R01 EB016089, R01 EB023963, R01 EB032788, R01 EB035529, R21 EB033516, K99 HD118185, R00 AG080084, and P41 EB031771.

## Notes

### Competing Interest Statement

The authors have declared no competing interest.

