## Supplementary material for "Correction for age-appropriate relaxation reference values alters estimates of neurometabolite concentration, but not age relationships"

| 1.Hardware | |
| --- | --- |
| a. Field strength [T] | 3 T |
| b. Manufacturer | Philips |
| c. Model (software version if available) | R 5.7.1 |
| d. RF coils: nuclei (transmit/receive), number of channels, type, body part | ^1^H, 32 channel, head |
| e. Additional hardware | - |
| 2. Acquisition | |
| a. Pulse sequence | PRESS |
| b. Volume of interest (VOI) locations | the CSO (predominantly WM) and PCC (predominantly GM) |
| c. Nominal VOI size [mm^3^] | 30 x 26 x 26 mm^3^ |
| d. Repetition time (TR), echo time (TE)[ms] | TR 2000 ms, TE 30 ms |
| e. Total number of averages per spectrum  i. Number of averaged specra per subspectrum | 1 total averages with 1 averages per subspectrum |
| f. Additional sequence parameters  i. Editing pulse parameters | F1: 2000 Hz, 2048 points |
| g. Water suppression method | VAPOR |
| h. Shimming method, reference peak, and threshold of acceptance of shim chosen | 1^st^ and 2^nd^ shimming pencil beam, water |
| i. Trigger or motion correction | No trigger or active motion correction |
| 3. Data analysis methods and outputs | |
| a. Analysis software | Osprey 2.9.0 |
| b. Processing steps deviating from Osprey | None |
| c. Output measure | tCr, rawWaterScaled, CSFWaterScaled, TissCorrWaterScaled, AlphaCorrWaterScaled, AlphaCorrWaterScaledGroupNormed |
| d. Quantification references and assumptions, fitting model assumptions | NAA, Cr, GPC, PCh, Lac, MMA, mI, Glu, Gln Asc, Asp, Cr, CrCH2, GABA, GPC, GSH, Gln, Glu, mI, Lac, NAA, NAAG, PCh, PE, sI, Tau, Lip09, Lip13, Lip20, MM_PRESS, tNAA, Glx, tCho, tCr.  Fitting method: Osprey baseline knot spacing 0.40 ppm |
| 4. Data quality | |
| PRESS, CSO – original analysis pipeline | |
| a. SNR (Cr), linewidth (Cr) [Hz, OFF spectra] | SNR 76, linewidth 7.05 Hz |
| b. Quality measures of postporcessing model fitting (Mean Relative Amplitude Residual (Residual/Noise)) | 3.87% |
| PRESS, PCC – original analysis pipeline | |
| a. SNR (Cr), linewidth (Cr) [Hz, OFF spectra] | SNR 93, linewidth 5.98 Hz |
| b. Quality measures of postporcessing model fitting (Mean Relative Amplitude Residual (Residual/Noise)) | 5.17% |
| PRESS, CSO – age-invariant pipeline | |
| a. SNR (Cr), linewidth (Cr) [Hz, OFF spectra] | SNR 93, linewidth 5.98 Hz |
| b. Quality measures of postporcessing model fitting (Mean Relative Amplitude Residual (Residual/Noise)) | 5.17% |
| PRESS, PCC – age-invariant pipeline | |
| a. SNR (Cr), linewidth (Cr) [Hz, OFF spectra] | SNR 102, linewidth 5.92 Hz |
| b. Quality measures of postporcessing model fitting (Mean Relative Amplitude Residual (Residual/Noise)) | 4.55% |
| PRESS, CSO – age-appropriate relaxation correction pipeline | |
| a. SNR (Cr), linewidth (Cr) [Hz, OFF spectra] | SNR 93, linewidth 5.98 Hz |
| b. Quality measures of postporcessing model fitting (Mean Relative Amplitude Residual (Residual/Noise)) | 5.17% |
| PRESS, PCC – age-appropriate relaxation correction pipeline | |
| a. SNR (Cr), linewidth (Cr) [Hz, OFF spectra] | SNR 102, linewidth 5.92 Hz |
| b. Quality measures of postporcessing model fitting (Mean Relative Amplitude Residual (Residual/Noise)) | 4.55% |

Supplementary Table 1: Summary following minimum reporting standards in MRS generated in Osprey See Lin et al. 'Minimum Reporting Standards for in vivo Magnetic Resonance Spectroscopy (MRSinMRS): Experts' consensus recommendations. NMR in Biomedicine. 2021;e4484. [doi.org/10.1002/nbm.4448](https://doi.org/10.1002/nbm.4448)
